# A GWAS-Derived Polygenic Score for Interleukin-1β is Associated with Hippocampal Volume in Two Samples

**DOI:** 10.1101/516989

**Authors:** Reut Avinun, Adam Nevo, Annchen R. Knodt, Maxwell L. Elliott, Ahmad R. Hariri

## Abstract

Accumulating research suggests that the pro-inflammatory cytokine interleukin-1β (IL-1β) has a modulatory effect on the hippocampus, a brain structure important for learning and memory as well as linked with both psychiatric and neurodegenerative disorders. Here, we use an imaging genetics strategy to test an association between an IL-1β polygenic score, derived from summary statistics of a recent genome-wide association study (GWAS) of circulating cytokines, and hippocampal volume, in two independent samples. In the first sample of 512 non-Hispanic Caucasian university students (274 women, mean age 19.78 ± 1.24 years) from the Duke Neurogenetics Study, we identified a significant positive correlation between higher polygenic scores, which presumably reflect higher circulating IL-1β levels, and average hippocampal volume. This positive association was successfully replicated in a second sample of 7,960 white British volunteers (4,158 women, mean age 62.63±7.45 years) from the UK Biobank. Collectively, our results suggest that a functional GWAS-derived score of IL-1β blood circulating levels affects hippocampal volume, and lend further support in humans, to the link between IL-1β and the structure of the hippocampus.

## Introduction

The hippocampus, a key brain structure supporting learning and memory, has been implicated in the pathophysiology of both psychiatric and neurodegenerative disorders. For example, smaller hippocampal volume has been noted in schizophrenia (van Erp et al., 2016), depression (Schmaal et al., 2016), and bipolar disorder (Hibar et al., 2016), as well as in neurodegenerative disorders such as Alzheimer’s disease (Bobinski et al., 1999). More broadly, smaller hippocampal volume has been associated with poorer cognitive function in healthy adults (Zhu et al., 2017). Thus, advancing the understanding of the factors that contribute to individual differences in hippocampal volume, can shed light on both normal and abnormal function.

It is now clear that inflammation has far-reaching modulatory effects on the central nervous system (Miller et al., 2013; Yirmiya and Goshen, 2011). Amongst these, the pro-inflammatory cytokine interleukin (IL)-1β may be particularly important for understanding hippocampal structure. First, receptors for IL-1β are highly expressed in the dentate gyrus (Ban et al., 1991; Loddick et al., 1998). Second, IL-1β has been shown to affect neurogenesis, synaptic strengthening, and long term potentiation (Avital et al., 2003; Bellinger et al., 1993; Goshen et al., 2008; Green and Nolan, 2012; Katsuki et al., 1990), which are all processes that can affect hippocampal volume. For example, in animal models, interrupting IL-1β signaling, by a deletion of IL-1 receptor type 1 or by administration of IL-1 receptor antagonist, blocks the antineurogenic effect of stress on the hippocampus. In contrast, increasing IL-1β signaling, by exogenous administration of this cytokine, mimics the effect of stress, and decreases hippocampal cell proliferation (Koo and Duman, 2008). In humans, Zunszain et al., (2012) demonstrated that IL-1β directly inhibits neurogenesis in hippocampal progenitor cells in vitro.

Despite the vast literature demonstrating the effects of IL-1β on hippocampal volume and studies showing that IL-1β levels are moderately to highly heritable (Brodin et al., 2015; De Craen et al., 2005), to our knowledge, the genetic influences of IL-1β on hippocampal volume in humans have only been previously shown in one candidate gene study (Raz et al., 2015). In a sample of 80 individuals, Raz et al. found that the T allele of the single nucleotide polymorphism (SNP) rs16944 in the IL-1β gene was associated with smaller hippocampal volume. Notably however, candidate gene studies suffer from low replicability rates (e.g., Avinun et al., 2018), and more specifically, findings regarding the biological functionality of rs16944 have not been consistent across studies (e.g., Chen et al., 2006; Iacoviello et al., 2005).

Polygenic scores that aggregate information from across the genome to summarize genome-wide genetic influences on an outcome of interest, have been successfully used in previous research to model individual differences (e.g., Domingue et al., 2015; Domingue et al., 2017; Stephan et al., 2018). A recent study conducted separate genome-wide analyses of circulating blood levels of 41 cytokines, including IL-1β, in up to 8,293 adult Finns from three independent population cohorts (Ahola-Olli et al., 2017). Here, we used summary statistics from this GWAS to generate polygenic scores to model potential individual differences in circulating levels of IL-1β. There are two main advantages to this approach: 1) whole genome data is more available to researchers than measures of IL-1β levels in blood and/or cerebrospinal fluid, and consequently findings with genetic scores can be more easily replicated and extended; and 2) the use of genetic data supports causal inference more than direct measures of IL-1β levels.

Based on the available literature, we hypothesized that higher polygenic IL-1β scores, which putatively index higher circulating levels of the pro-inflammatory cytokine, would be associated with smaller hippocampal volume. We tested our hypothesis in two independent samples: 1) 512 non-Hispanic Caucasian university students from the Duke Neurogenetics Study; and 2) 7,960 adult white British volunteers from the UK Biobank.

## Materials and methods

### Participants

Our first sample consisted of 512 self-reported non-Hispanic Caucasian participants (274 women, mean age 19.78±1.24 years) from the Duke Neurogenetics Study (DNS) for whom there was complete data on genotypes, structural MRI, and all covariates detailed below. All procedures were approved by the Duke University Medical Center Institutional Review Board, and participants provided informed written consent before study initiation. All participants were free of the following study exclusions: 1) medical diagnoses of cancer, stroke, diabetes requiring insulin treatment, chronic kidney or liver disease, or lifetime history of psychotic symptoms; 2) use of psychotropic, glucocorticoid, or hypolipidemic medication; and 3) conditions affecting cerebral blood flow and metabolism (e.g., hypertension).

Current and lifetime DSM-IV Axis I or select Axis II disorders (antisocial personality disorder and borderline personality disorder), were assessed with the electronic Mini International Neuropsychiatric Interview (Lecrubier et al., 1997) and Structured Clinical Interview for the DSM-IV Axis II subtests (First et al., 1997), respectively. Of the 512 participants with data included in our analyses, 114 individuals had at least one DSM-IV diagnosis. Importantly, neither current nor lifetime diagnosis were an exclusion criterion, as the DNS seeks to establish broad variability in multiple behavioral phenotypes related to psychopathology. However, no participants, regardless of diagnosis, were taking any psychoactive medication during or at least 14 days prior to their participation.

Our second sample, consisted of 7,960 white British participants (4,158 women, mean age 62.63±7.45 years), with complete genotype, structural MRI, and covariate data from the UK Biobank (www.ukbiobank.ac.uk; Sudlow et al., 2015), which includes over 500,000 participants, between the ages of 40 and 69 years, who were recruited within the UK between 2006 and 2010. The UK Biobank study has been approved by the National Health Service Research Ethics Service (reference: 11/NW/0382), and our analyses were conducted under UK Biobank application 28174.

### Socioeconomic status (SES)

Prior research has reported associations between SES and brain volume (Hackman et al., 2010). In the DNS, we controlled for possible SES effects using the “social ladder” instrument (Adler et al., 2000), which asks participants to rank themselves relative to other people in the United States (or their origin country) on a scale from 0—10, with people who are best off in terms of money, education, and respected jobs, at the top (10) and people who are worst off at the bottom (0).

In the UK Biobank, we used the Townsend deprivation index, which was assessed at recruitment, as a means to control for SES. The Townsend deprivation index is a composite measure of deprivation based on unemployment, non-car ownership, non-home ownership, and household overcrowding. It was calculated before participants joined the UK Biobank and was based on the preceding national census data, with each participant assigned a score corresponding to the postcode of their home dwelling. Scoring was reversed so that high values represented high SES.

### Body mass index (BMI)

Prior research has reported associations between BMI and brain volume (Gunstad et al., 2008). In both DNS and UK Biobank samples, BMI was calculated based on the height and weight of the participants. In the DNS, this calculation was based on imperial system values (pounds/inches *703), while in the UK Biobank the metric system was used (kg/m^2^). For the UK Biobank we used the BMI from the imaging assessment visit.

### Recent life stress

Prior research has reported associations between stress and hippocampal volume (Lupien et al., 2009). In the DNS, we controlled for the effects of life stress during the year prior to assessment using a summation of 38 negatively valenced items (as described in Avinun et al., 2017; Nikolova et al., 2012) from the Life Events Scale for Students (LESS; Clements and Turpin, 1996).

In the UK Biobank, life stress during the two years prior to imaging was assessed based on a count of 6 stressful events (illness of participant, illness of a close relative, death of a partner/spouse, death of a close relative, marital separation/divorce, and financial difficulties).

### Race/Ethnicity

Because self-reported race and ethnicity are not always an accurate reflection of genetic ancestry, an analysis of identity by state of whole-genome SNPs was performed in PLINK (Purcell et al., 2007). The first two multidimensional scaling components within the non-Hispanic Caucasian subgroup were used as covariates in analyses of data from the DNS. The decision to use only the first two components was based on an examination of a scree plot of the variance explained by each component.

For analyses of data from the UK Biobank, only those who were ‘white British’ based on both self-identification and a principal components analysis of genetic ancestry were included. Additionally, the first 10 multidimensional scaling components received from the UK biobank’s team were included as covariates as previously done (e.g., Whalley et al., 2016).

### Genotyping

In the DNS, DNA was isolated from saliva using Oragene DNA self-collection kits (DNA Genotek) customized for 23andMe (www.23andme.com). DNA extraction and genotyping were performed through 23andMe by the National Genetics Institute (NGI), a CLIA-certified clinical laboratory and subsidiary of Laboratory Corporation of America. One of two different Illumina arrays with custom content was used to provide genome-wide SNP data, the HumanOmniExpress (N=326) or HumanOmniExpress-24 (N=186; Do et al., 2011; Eriksson et al., 2010; Tung et al., 2011).

In the UK Biobank, samples were genotyped using either the UK BiLEVE (N=775) or the UK Biobank axiom (N=7,185) array. Details regarding the UK Biobank’s quality control can be found elsewhere (Bycroft et al., 2017).

### Quality control and polygenic scoring

For genetic data from both the DNS and UK Bionbank, PLINK v1.90 (Purcell et al., 2007) was used to perform quality control analyses and exclude SNPs or individuals based on the following criteria: missing genotype rate per individual >.10, missing rate per SNP >.10, minor allele frequency <.01, and Hardy-Weinberg equilibrium p<1e-6. Additionally, in the UK Biobank, quality control variables that were provided with the dataset were used to exclude participants based on a sex mismatch (genetic sex different from reported sex), a genetic relatedness to another participant, and outliers for heterozygosity or missingness.

Polygenic scores were calculated using PLINK’s (Purcell et al., 2007) “--score” command based on published SNP-level summary statistics from a recent GWAS that included IL-1β blood levels as an outcome of interest (Ahola-Olli et al., 2017). SNPs from the IL-1β GWAS were matched with SNPs from the DNS and UK Biobank datasets. For each SNP the number of the alleles (0, 1, or 2) associated with IL-1β blood levels was multiplied by the effect estimated in the GWAS. The polygenic score for each individual was an average of weighted IL-1β-associated alleles. All SNPs that could be matched with SNPs from the DNS or UK biobank were used regardless of effect size and significance in the original GWAS, as previously recommended and shown to be effective (Dudbridge, 2013; Ware et al., 2017). A total of 441,939 SNPs in the DNS and 645,022 SNPs in the UK Biobank were included in the respective polygenic scores. The approach described here for the calculation of the polygenic score was successfully used in previous studies (e.g., Domingue et al., 2015; Domingue et al., 2017; Stephan et al., 2018).

### Structural MRI

In the DNS, data were collected at the Duke-UNC Brain Imaging and Analysis Center using one of two identical research-dedicated GE MR750 3T scanners (General Electric Healthcare, Little Chalfont, United Kingdom) equipped with high-power high-duty cycle 50-mT/m gradients at 200 T/m/s slew rate, and an eight-channel head coil for parallel imaging at high bandwidth up to 1 MHz. T1-weighted images were obtained using a 3D Ax FSPGR BRAVO with the following parameters: TR = 8.148 ms; TE = 3.22 ms; 162 axial slices; flip angle, 12°; FOV, 240 mm; matrix =256×256; slice thickness = 1 mm with no gap; and total scan time = 4 min and 13 s.

To generate regional measures of brain volume, anatomical images for each subject were first skull-stripped using ANTs (Klein et al., 2009), then submitted to Freesurfer’s (Version 5.3) recon-all with the “-noskullstrip” option (Dale et al., 1999; Fischl et al., 1999), using an x86_64 linux cluster running Scientific Linux. The gray and white matter boundaries determined by recon-all were visually inspected using FreeSurfer QA Tools (https://surfer.nmr.mgh.harvard.edu/fswiki/QATools) and determined to be sufficiently accurate for all subjects. Volume measures for the hippocampus from each participant’s aseg.stats file were averaged across hemispheres. Estimated Total Intracranial Volume (eTIV) was used to quantify intracranial volume (ICV).

In the UK Biobank, imaging data were collected on a Siemens Skyra 3T, with a standard Siemens 32-channel RF receive head coil, and preprocessed with FSL packages (the FMRIB Software Library; Jenkinson et al., 2012). Segmentation of T1-weighted structural images into subcortical structures was done using FIRST (FMRIB’s Integrated Registration and Segmentation Tool; Patenaude et al., 2011). Here as well, left and right hippocampal volumes were averaged to create a mean volume variable. ICV was estimated based on the sum of white matter, gray matter and ventricular cerebrospinal fluid volumes. Further details for the UK Biobank imaging protocol can be found at http://biobank.ctsu.ox.ac.uk/crystal/refer.cgi?id=1977.

### Statistical analyses

Mplus version 7 (Muthén and Muthén, 2007) was used to conduct linear regression analyses. In both samples the covariates of no interest included: participants’ sex (coded as 0=males, 1=females), age (in the DNS 18-22 years were coded as 1-5), ethnicity components, BMI, SES, recent life stress, and intracranial volume. The IL-1β score, intracranial volume, and hippocampal volume were standardized to improve interpretability. Maximum likelihood estimation with robust standard errors, which is robust to non-normality, was used in the regression analyses. Standardized results are presented.

## Results

Descriptive statistics for study variables and correlations are presented in Table 1 and Table 2, respectively. Interestingly, the IL-1β score was positively associated with SES in both samples, therefore the findings below are reported with and without SES as a covariate to avoid multicollinearity. The polygenic scores did not differ between men and women (DNS: F[1, 510] = .642, p=.42; UK Biobank: [1, 7958] = .021, p=.88).

**Table 1.**
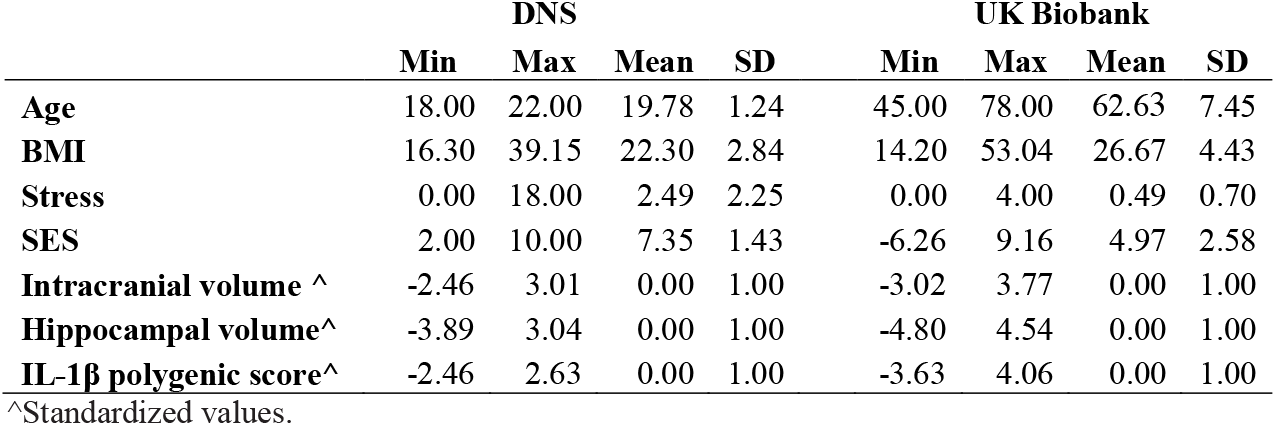
Descriptive statistics of study variables.

**Table 2.** Correlations between study variables.

|  | DNS |  |  |  |  |  | UK Biobank |  |  |  |  |  |
| --- | --- | --- | --- | --- | --- | --- | --- | --- | --- | --- | --- | --- |
|  | IL-1B<br>polygenic<br>score | Age | SES | Recent<br>life<br>stress | BMI | ICV | IL-1B<br>polygenic<br>score | Age | SES | Recent<br>life<br>stress | BMI | ICV |
| **IL-1B polygenic score** | 1 |  |  |  |  |  | 1 |  |  |  |  |  |
| **Age** | -.008 | 1 |  |  |  |  | -.021^ | 1 |  |  |  |  |
| **SES** | .099* | .115** | 1 |  |  |  | .024* | .083** | 1 |  |  |  |
| **Recent life stress** | -0.038 | -.091* | -.125** | 1 |  |  | 0 | -.147** | -.068** | 1 |  |  |
| **BMI** | -0.077^ | .022 | -0.079^ | 0.017 | 1 |  | -0.011 | -.031** | -0.071** | .077** | 1 |  |
| **ICV** | 0.03 | -.01 | .114* | -0.037 | .115** | 1 | 0.007 | -.163** | 0.026* | -0.026* | .067** | 1 |
| **Hippocampal volume** | .097* | 0.017 | .122** | -0.016 | 0.075 | .645** | .030** | -.241** | 0.011 | -0.012 | .023* | .466** |

^p<.10, \*p<.05, \*\*p<.01

The Analysis in the DNS sample revealed that IL-1β polygenic scores significantly predicted hippocampal volume, so that scores that were associated with higher IL-1β levels, were associated with larger volume (without SES: β=.078, SE=.037, p=.036; with SES: β=.075, SE=.037, p=.044). Of the covariates, only ICV was significantly associated with hippocampal volume (β=.626, SE=.044, p<.001). In the UK Biobank analysis, IL-1β polygenic score was also positively and significantly associated with hippocampal volume (without SES: β=.023, SE=.010, p=.020; with SES: β=.022, SE=.010, p=.021). Of the covariates, other than ICV (β=.425, SE=.014, p<.001), both age (β=−.177, SE=.010, p<.001) and stress (β=−.025, SE=.010, p=.010) negatively predicted hippocampal volume.

As post-hoc analyses we tested each hemisphere separately with the same covariates, including SES. In the DNS the effect was stronger and only significant in the left hippocampus (LEFT: β=.109, SE=.039, p=.005; RIGHT: β=.024, SE=.036, p=.503). In the UK biobank, the effect was also only significant in the left hippocampus (LEFT: β=.022, SE=.010, p=.025; RIGHT: β=.018, SE=.010, p=.072).

Additionally, in the DNS we were able to test whether a DSM-IV diagnosis, as indicated by a clinical interview, affected the findings. With the addition of a variable indicating a DSM-IV diagnosis (0-no diagnosis; 1-at least one DSM-IV diagnosis) as a covariate, the positive association of the IL-1β polygenic score with hippocampal volume remained significant (β=.074, SE=.037, p=.047).

## Discussion

Here we report that a polygenic score for circulating levels of the pro-inflammatory cytokine IL-1β, based on summary statistics from a recent GWAS, is associated with hippocampal volume in two independent samples: the DNS, which consists of high-functioning 18-22 year old university students, and the UK biobank, which consists of 45-78 year old volunteers. Contrary to our hypothesis, which was based on previous research demonstrating that high IL-1β levels suppress neurogenesis (Goshen et al., 2008; Koo and Duman, 2008; Zunszain et al., 2012), we found that higher IL-1β polygenic scores, which putatively index increased levels of this pro-inflammatory cytokine, were associated in both samples with larger hippocampal volume.

While IL-1β may adversely affect neurogenesis, it has been shown to increase gliogenesis (Chen et al., 2013; Crampton et al., 2012). In other words, IL-1β has been shown to reduce neurogenesis, not by increasing cell death, but by affecting cell fate determination and promoting the differentiation of neural progenitor cells into glial cells (Crampton et al., 2012). Additional processes that may explain the observed positive association between IL-1β polygenic scores and hippocampal volume, are the effects of IL-1β on long term potentiation (LTP; Avital et al., 2003) and dendritic spine size (Goshen et al., 2009). Mice with impaired IL-1 signaling are characterized by deficits in memory function, reduced LTP, and reduced dendritic spine size (Avital et al., 2003; Goshen et al., 2008; Goshen et al., 2007; Yirmiya et al., 2002), suggesting that certain levels of IL-1β are required for normal brain function. It is possible that the higher levels of IL-1β, as modeled by the polygenic score, are within an optimal range that leads to an advantageous function of this cytokine in promoting synaptic integrity and plasticity. Further studies, combining genetic, immune, and structural brain measurements during various stages in development are needed to shed light on the processes that may underlie the positive association between IL-1β polygenic scores and hippocampal volume.

In addition to our primary analyses of IL-1β polygenic scores and hippocampal volume, we observed a weak positive correlation between IL-1β scores and SES in both the DNS and UK Biobank samples. This positive correlation is also surprising, as lower SES has been associated with higher levels of inflammation (Loucks et al., 2010). Of note, this association is usually interpreted as an effect of SES on inflammation. However, our current observation suggests the opposite direction may also be possible (i.e., inflammation affecting SES), and may be of interest to future research on SES and environmental risk. As noted above, however, we have no data on circulating cytokines and the directionality of these links is speculative.

Although our study has several strengths, including the use of two independent samples with markedly different characteristics (e.g., young university students versus older community volunteers) and a GWAS-derived functionally informed polygenic score, it is not without limitations. First, our analyses and findings are restricted to non-Hispanic Caucasians. Future studies are necessary to determine whether the IL-1β polygenic score can be generalized to other racial and ethnic populations. Second, we did not examine specific hippocampal subregions, such as the dentatge gyrus, which may exhibit preferential sensitivity to variability in IL-1β signaling (Ban et al., 1991; Loddick et al., 1998). Such subregional analyses require higher resolution structural data. Lastly, as cytokine levels were not measured in our samples, we were not able to validate the functionality of the IL-1β score or examine how it correlates with other pro-inflammatory markers. These limitations notwithstanding, our results further support the association between IL-1β levels and hippocampal volume in humans and motivate additional research on the links between the IL-1β polygenic score, inflammation, and brain volume.

## ACKNOWLEDGEMENTS AND DISCLOSURES

We thank the Duke Neurogenetics Study participants, the staff of the Laboratory of NeuroGenetics, and Emily Drabant Conley of 23andMe, Inc. We would also like to thank Dr. Megan Cooke for commenting on an early draft of the manuscript. The Duke Neurogenetics Study received support from Duke University as well as US-National Institutes of Health grants R01DA033369 and R01DA031579. RA, ARK, and ARH received further support from US-National Institutes of Health grant R01AG049789. MLE was supported by the National Science Foundation Graduate Research Fellowship under Grant No. NSF DGE-1644868. The Brain Imaging and Analysis Center was supported by the Office of the Director, National Institutes of Health under Award Number S10OD021480. Lastly, we would like to thank the UK Biobank participants and the UK Biobank team.

## Author contributions

RA, AN, and AH designed research; AK, and AH performed research; RA, AK, and ME analyzed data; RA, AN, AK, ME, and AH, wrote the paper.

